# Identification of Growth Differentiation Factor-15 as An Early Predictive Biomarker for Metabolic Dysfunction-Associated Steatohepatitis: A Nested Case-control Study of UK Biobank Proteomic Data

**DOI:** 10.1101/2024.09.12.612665

**Authors:** Hao Wang, Xiaoqian Xu, Yameng Sun, Hong You, Jidong Jia, You-Wen He, Yuanyuan Kong

**Author notes:** **Correspondence to:** Prof. Yuanyuan Kong, PhD, National Clinical Research Center for Digestive Diseases, State Key Lab of Digestive Health, Beijing Friendship Hospital, Capital Medical University; Clinical Epidemiology and EBM Unit, Beijing Clinical Research Institute, Beijing, China.

## Abstract

**Background & Aims:** The lack of non-invasive biomarkers for the early prediction of patients with metabolic dysfunction-associated steatohepatitis (MASH) is a major challenge for timely intervention. This study aims to determine the predictive capability for MASH long before its diagnosis by using six previously identified diagnostic biomarkers for metabolic dysfunction-associated steatotic liver disease (MASLD) with proteomic data from the UK Biobank.

**Methods:** A nested case-control study comprising of a MASH group and three age- and sex-matched controls groups (metabolic dysfunction-associated steatosis, viral hepatitis, and normal liver controls) were conducted. Olink proteomics, anthropometric and biochemical data at baseline levels were obtained from the UK Biobank. The baseline levels of CDCP1, FABP4, FGF21, GDF15, IL-6 and THBS2 were analyzed prospectively to determine their predictive accuracy for subsequent diagnosis with a mean lag time of over 10 years.

**Results:** At baseline, GDF15 demonstrated the best performance for predicting MASH occurrence at 5 and 10 years later, with an AUC of 0.90 at 5 years and 0.86 at 10 years. A predictive model based on four biomarkers (GDF15, FGF21, IL-6, and THBS2) showed AUCs of 0.88 at both 5 and 10 years. Furthermore, a protein-clinical model that included these four circulating protein biomarkers along with three clinical factors (BMI, ALT and TC) yielded AUCs of 0.92 at 5 years and 0.89 at 10 years.

**Conclusion:** GDF15 at baseline levels outperformed other individual circulating protein biomarkers for the early prediction of MASH. Our data suggest that GDF15 and the GDF15-based model may be used as easy-to-implement tools to identify patients with high risk of developing MASH at a mean lag time of over 10 years.

## Introduction

Metabolic dysfunction-associated steatotic liver disease (MASLD), formerly known as non-alcoholic fatty liver disease (NAFLD), affects about 38% of the global population and is the most common chronic liver disease worldwide.(1) MASLD encompasses a spectrum of disease severity, including hepatic steatosis with or without mild inflammation (metabolic dysfunction-associated steatosis, MAS) and a necroinflammatory subtype (metabolic dysfunction-associated steatohepatitis, MASH). MASH, the active form of MASLD, is characterized by the histological lobular inflammation, hepatocyte ballooning and faster fibrosis progression.(2) The global prevalence of MASH is 5.27%,(3) and it has become the second or third leading cause of end-stage liver disease and hepatocellular carcinoma (HCC).(4, 5) A large proportion of MASH patients remain undiagnosed until the disease has progressed to more advanced and life-threatening stages due to its largely asymptomatic nature in the early stage.(6) Thus, early prediction of patients at high risk of MASH is an important medical need and will play a significant role in the timely intervention and personalized treatment from the perspective of preventive medicine.

To date, no biomarker is available to predict MASH development. Histological assessment of liver biopsy remains the gold standard for diagnosing MASH and staging liver fibrosis. However, the limitations of biopsy, including its invasiveness, associated complications, high cost, time-consuming nature and intra-observer variability, make it less suitable for MASH monitoring in clinical practice, and unsuitable for screening in the population-based health surveys.(7) Imaging based diagnostic tools, such as ultrasonography and magnetic resonance imaging, are primarily used for detecting MASH and fibrosis, but these tests cannot reliably differentiate patients with MASH from patients with MAS and lack predictive capability for MASH disease development.(8) Therefore, developing innovative and non-invasive biomarkers for predicting MASH holds a promising prospect for clinical application.

During the last decade, six non-invasive protein biomarkers have been identified as diagnostic biomarkers to differentiate patients with MASH, MASLD, and advanced fibrosis.(9, 10) These biomarkers include CUB domain-containing protein 1 (CDCP1), fatty acid binding protein 4 (FABP4), fibroblast growth factor 21 (FGF21), growth differentiation factor 15 (GDF15), interleukin-6 (IL-6), and thrombospondin 2 (THBS2). CDCP1 is a transmembrane glycoprotein involved in the tyrosine phosphorylation-dependent regulation of cellular events related to tumor invasion and metastasis.(11) Recent omics-based studies have identified serum soluble CDCP1 as a robust serological biomarker for risk stratification of patients with MASH.(12) FABP4 is a fatty acid binding protein that participates in fatty acid uptake, transport, and metabolism.(13) A previous study found that FABP4 level could be a potential biomarker for MASLD diagnosis.(14) FGF21 is a stress-inducible hormone that has important roles in regulating energy balance and glucose and lipid homeostasis. FGF21 is a potential pharmaceutical target for obesity-related metabolic complications, including hyperglycemia, dyslipidemia, and MASH.(15) GDF15 is a member of the TGF-β superfamily whose expression increases in response to cellular stress and diseases.(16) GDF-15 levels rise with the progression of MASLD severity.(17) IL-6 is a cytokine that functions in inflammation and regulation of metabolic homeostasis.(18) Increasing evidence indicates that IL-6 is involved in metabolic diseases, including MASLD and MASH.(19, 20) THBS2 is a homotrimeric glycoprotein mediating cell-to-cell and cell-to-matrix interactions, and a key determinant of fibrogenesis in MASLD.(21) These identified diagnostic biomarkers have shown potential in distinguishing between different stages and severities of liver disease, highlighting their significance in the early prediction and management of MASH.

Based on the identification of the above six circulating diagnostic biomarkers for MASLD and MASH, we hypothesize that these soluble non-invasive biomarkers may serve as predictive biomarkers for MASH development long before disease diagnosis. To test this hypothesis, we analyzed the proteomic data from the prospective UK Biobank study which recruits human subjects, establishes their multi-omics at baseline levels, and then follows disease development in the entire cohort. Here, we report the identification of GDF-15 as a predictive biomarker for MASH development long before disease diagnosis.

## Materials and Methods

### Study design

To validate proteomic biomarkers within the UK Biobank, a nested case-control study was conducted. This approach involves selecting cases and controls from the population of an existing cohort, offering several advantages over traditional case-control studies. Firstly, this method allows for the selection of cases and controls from the same population, which enhances internal validity by ensuring that both groups are drawn from the same underlying cohort and demographic characteristics. This is crucial in minimizing biases that can arise from differences in population characteristics. Secondly, nested case-control studies often employ matching techniques to pair cases with controls based on factors such as age, sex, and other relevant variables. This matching helps to minimize potential confounding variables, thereby making the analysis of biomarkers more robust and reliable. Overall, the nested case-control design within the UK Biobank provides a powerful framework for validating proteomic biomarkers, ensuring rigorous evaluation and enhancing the potential for meaningful insights into disease mechanisms and predictive biomarker performance.

The study was approved by the North West Multi-Centre Research Ethics Committee, and detailed information about its methodology and protocols can be found on the official UK Biobank website (http://www.ukbiobank.ac.uk). The current study’s protocol is also available online, conducted under the UK Biobank application number 129053. This large-scale initiative facilitates extensive research into various health conditions by linking comprehensive baseline data with long-term health outcomes.

### Definition of cases and multiple controls

In the study conducted using data from the UK Biobank and based on the International Classification of Diseases, 10th revision (ICD-10), four distinct groups were identified from hospital medical records: MASH, MAS, viral hepatitis, and normal liver groups. Below is how each group was defined and selected for comparison:

#### MASH group

Participants diagnosed with MASH were defined as the case group. Consistent with the Expert Panel Consensus statement, MASH cases were defined as ICD-10 K75.8 and were excluded if they had: (1) a prior diagnosis of MASH/MASLD before blood sample collection; (2) a diagnosis of MASH/MASLD within six months from recruitment; or (3) a diagnosis of other liver diseases (detailed in Supplemental Table 1).

#### Control groups

Three control groups were established for comprehensive comparison: MAS, viral hepatitis, and normal liver groups. MAS group was defined as ICD-10 K76.0 and were excluded if they had: (1) a prior diagnosis of MASH/MASLD before blood sample collection; (2) a diagnosis of MASH/MASLD within six months from recruitment; or a (3) diagnosis of other liver diseases (**Table S1**). Virial hepatitis group was defined as ICD-10 B15.0, B16.0, B17.0, B18.0, B19.0, and were excluded if they had: (1) a prior diagnosis of virial hepatitis before blood sample collection; or (2) a diagnosis of other liver diseases. Participants without any medical records of liver diseases were defined as normal liver controls.

#### Matching Protocol

MASH cases and non-MASH controls were matched on age (within the same year) and sex in a 1:2 ratio to minimize confounding factors. However, due to sample size constraints, viral hepatitis controls were matched with MASH cases in a 1:1 ratio. Non-MASH controls were randomly selected from their respective groups to ensure unbiased comparison. This matching and selection strategy aimed to elucidate the unique proteomic profiles associated with each condition (MASH, MAS, viral hepatitis, and normal liver), facilitating a comprehensive analysis of biomarkers and their potential diagnostic utility in differentiating these liver disease states within the UK Biobank cohort.

### Anthropometric and Biochemical Measurements

Age at baseline was determined from the participants’ dates of birth and the dates of their baseline assessments. Sex was self-reported at baseline. Anthropometric measurements, including height and weight, were collected. Body mass index (BMI) was calculated as weight in kilograms divided by height in meters squared (kg/m^2^).

Routine biochemical tests were conducted to measure of the levels of several biomarkers, including alanine transaminase (ALT), aspartate transaminase (AST), albumin (ALB), alkaline phosphatase (ALP), gamma-glutamyl transferase (GGT), platelet (PLT), total bilirubin (TBiL), total cholesterol (TC), triglycerides (TG), high-density lipoprotein (HDL), low-density lipoprotein (LDL), uric acid (UA), urea, C-reactive protein (CRP), and fasting plasma glucose (FPG). These measurements were obtained from the UK Biobank and provide a comprehensive biochemical profile for each participant. This data, along with the proteomic data and anthropometric measurements, was used to analyze and validate the six candidate protein biomarkers (CDCP1, FABP4, FGF21, GDF15, IL-6, and THBS2) for their predictive capability in identifying individuals at high risk of developing MASH.

### Statistical analysis

Continuous variables were presented as means and standard deviations (SD) if normally distributed, or medians and interquartile ranges if not. Differences among case and control groups were tested using one-way ANOVA for normal data, or the Kruskal-Wallis test for non-normal data. For comparisons between two groups, the Student’s t-test was used for normal data, and the Mann-Whitney *U*-test for non-normal data. Categorical variables were presented as frequencies and percentages and analysed using the Chi-square test or Fisher’s exact test. To adjust for multiple comparisons, the Bonferroni correction was applied, with the adjusted alpha level calculated as αadjust=α_0_/κ (22). To identify potential proteomic biomarkers for predicting MASH, the expression of candidate proteomic biomarkers at baseline among MASH patients and control groups was illustrated using violin plots. Spearman’s rank correlation was employed to estimate the associations between candidate proteomic biomarkers and metabolic parameters (e.g., FPG, TC, TG) and liver function tests (e.g., ALT, AST, ALB) of the participants.

Multivariable Cox proportional hazards regression analyses were performed to identify independent proteomic biomarkers for MASH, adjusting for three different models. Hazard ratios (HRs) and 95% confidence intervals (CIs) were calculated to estimate the risk effects of candidate proteomic biomarkers for MASH. The proteomic biomarkers adjusting for different models were graphed by forest plot, which was generated using the R package ‘‘forestplot.’’ To further improve the predictive accuracy of potential protein biomarkers for MASH, Cox regression models were performed to construct predictive models based on the candidate protein biomarkers and MASH-related clinical risk factors. The time-dependent receiver operating characteristic (ROC) curves, which measure the performance of candidate biomarkers given the disease status of individuals at certain time points, were used to evaluate the predictive performance of the models. The area under the curve (AUC) and 95% bootstrap CI were calculated to assess the predictive performance of the models. The performance of the predictive models was evaluated using 10-fold cross-validation on the training set.

Statistical analyses were performed using R, version 4.4.0 (R Foundation for Statistical Computing) and SPSS 25.0 (IBM Corporation, New York, USA). Two-tailed P < 0.05 was considered statistically significant. Additional information on UK Biobank study population and Olink proteomics is included in Supplemental Methods.

## Results

### Baseline characteristics of the participants

A total of 53,014 participants with Olink proteome data were obtained from UK Biobank. According to their ICD-10 codes, 82 MASH cases, 831 MAS cases, 114 viral hepatitis cases and 51,987 normal liver healthy controls were identified. Among the 82 MASH cases, four patients with MASH at baseline level were excluded from this study. Among the 831 MAS cases, 68 patients with MAS at baseline level, 9 patients with other concurrent liver diseases, and 7 patients diagnosed with MAS within six months were excluded from this study. Based on the matching criteria, a total of 78 MASH cases and three groups of age- and sex matched controls, including 156 MAS cases, 78 viral hepatitis cases, and 156 normal liver healthy controls, were included in this study. This selection process ensures that the study groups are well-matched and minimizes potential confounding factors, thereby improving the reliability of the comparative analyses (**Fig. 1A**).

**Fig. 1.**
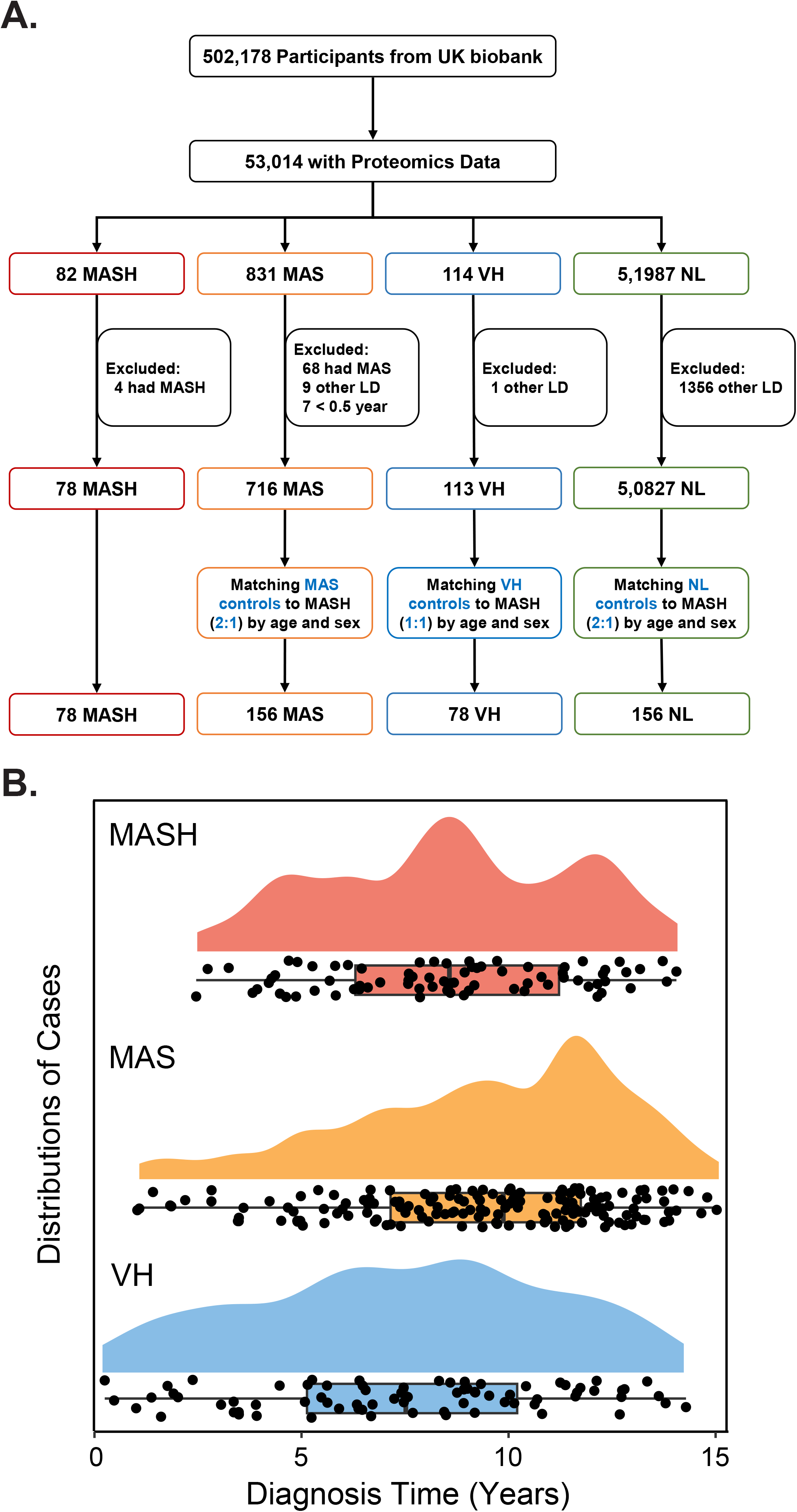
Flow chart of participant selection and mean time to diagnosis of liver diseases. (**A**) A total of 78 cases of MASH case were selected from the prospective cohort of 502,178 participants from an initial 82 cases. MASH, metabolic dysfunction-associated steatohepatitis; MAS, metabolic dysfunction-associated steatosis, defined as MASLD without steatohepatitis. VH: viral hepatitis. NL: normal liver function control subjects. (B) Distributions of diagnosis time for individual patients. Diagnosis time was defined as the lag time from recruitment (baseline) to disease diagnosis. Each black dot represents a single patient. Box represents mean and SD

The mean times to diagnosis for MASH, MAS and viral hepatitis cases are 12.93, 11.59, and 12.63 years, respectively (**Fig. 1B**). The demographic, anthropometric, and biochemical characteristics of participants are described in **Table 1**. The mean ages of MASH, MAS, viral hepatitis, and normal liver control groups were approximately 59 years, with each group comprising 40 males (52.63%) and 38 females (52.63%).

**Table 1.**
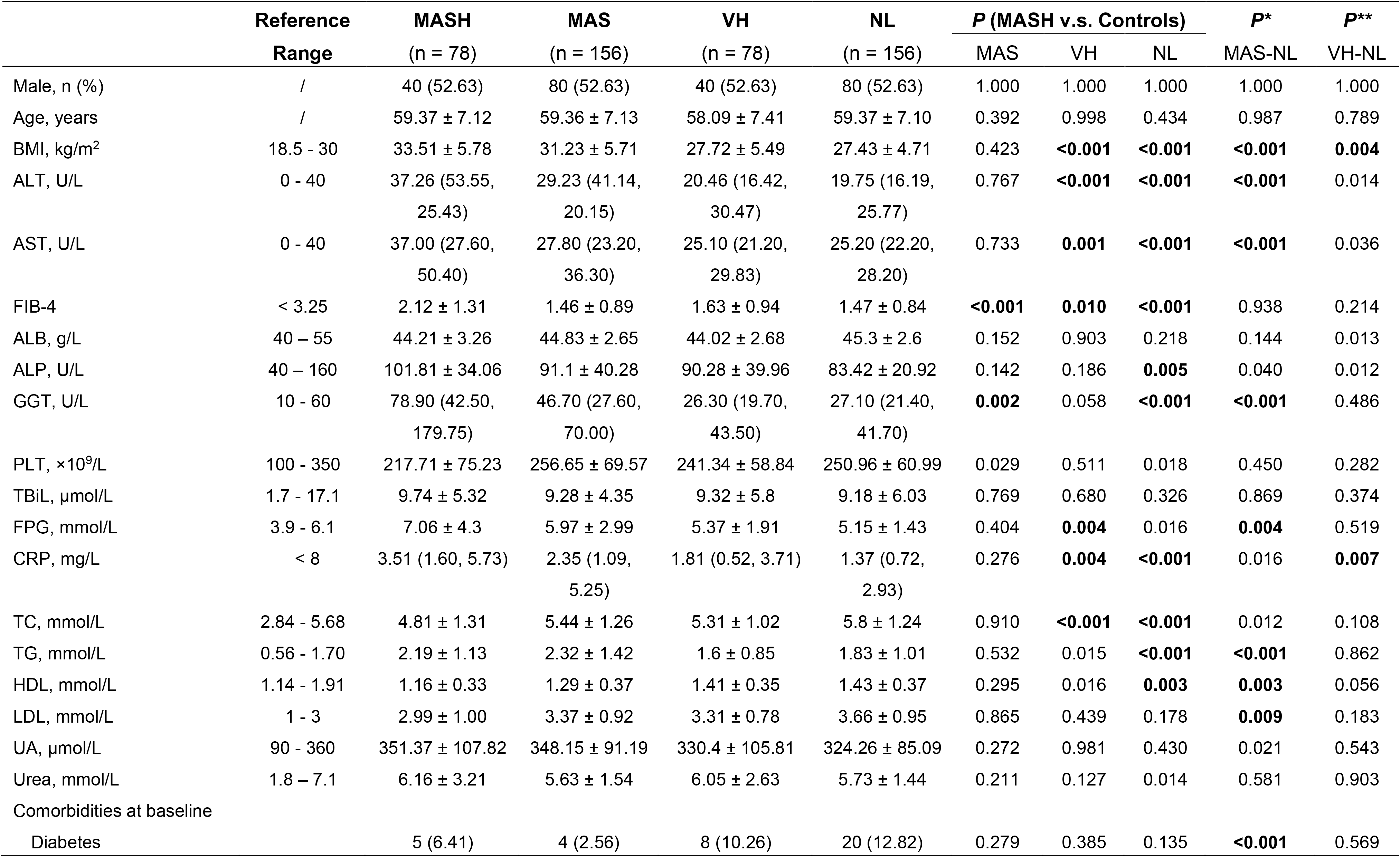
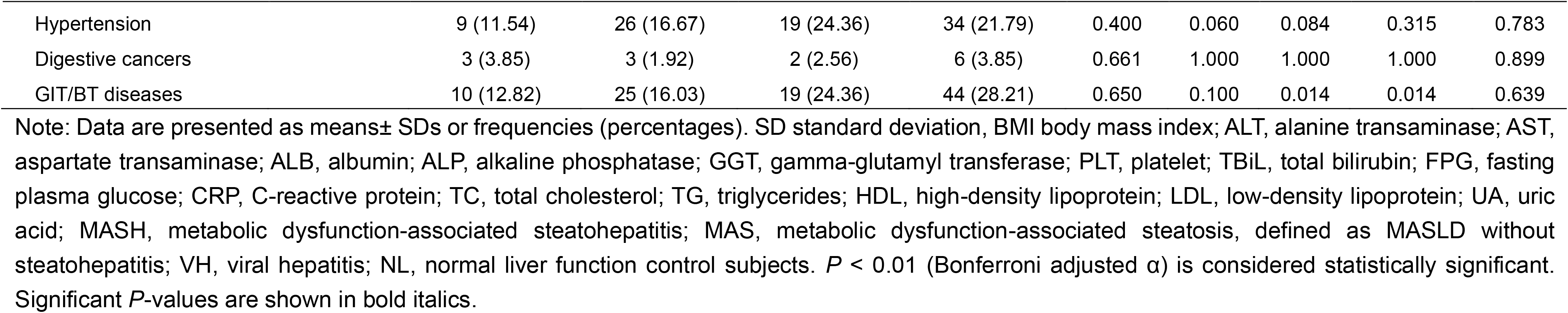
Clinical characteristics of the participants.

|  | Reference<br>Range | MASH<br>(n = 78) | MAS<br>(n = 156) | VH<br>(n = 78) | NL<br>(n = 156) | P (MASH v.s. Controls) |  |  | P* | P** |
| --- | --- | --- | --- | --- | --- | --- | --- | --- | --- | --- |
|  |  |  |  |  |  | MAS | VH | NL |  |  |
| Male, n (%) | / | 40 (52.63) | 80 (52.63) | 40 (52.63) | 80 (52.63) | 1.000 | 1.000 | 1.000 | 1.000 | 1.000 |
| Age, years | / | 59.37 ± 7.12 | 59.36 ± 7.13 | 58.09 ± 7.41 | 59.37 ± 7.10 | 0.392 | 0.998 | 0.434 | 0.987 | 0.789 |
| BMI, kg/m<sup>2</sup> | 18.5 - 30 | 33.51 ± 5.78 | 31.23 ± 5.71 | 27.72 ± 5.49 | 27.43 ± 4.71 | 0.423 | **<0.001** | **<0.001** | **<0.001** | **0.004** |
| ALT, U/L | 0 - 40 | 37.26 (53.55,<br>25.43) | 29.23 (41.14,<br>20.15) | 20.46 (16.42,<br>30.47) | 19.75 (16.19,<br>25.77) | 0.767 | **<0.001** | **<0.001** | **<0.001** | 0.014 |
| AST, U/L | 0 - 40 | 37.00 (27.60,<br>50.40) | 27.80 (23.20,<br>36.30) | 25.10 (21.20,<br>29.83) | 25.20 (22.20,<br>28.20) | 0.733 | **0.001** | **<0.001** | **<0.001** | 0.036 |
| FIB-4 | < 3.25 | 2.12 ± 1.31 | 1.46 ± 0.89 | 1.63 ± 0.94 | 1.47 ± 0.84 | **<0.001** | **0.010** | **<0.001** | 0.938 | 0.214 |
| ALB, g/L | 40 – 55 | 44.21 ± 3.26 | 44.83 ± 2.65 | 44.02 ± 2.68 | 45.3 ± 2.6 | 0.152 | 0.903 | 0.218 | 0.144 | 0.013 |
| ALP, U/L | 40 – 160 | 101.81 ± 34.06 | 91.1 ± 40.28 | 90.28 ± 39.96 | 83.42 ± 20.92 | 0.142 | 0.186 | **0.005** | 0.040 | 0.012 |
| GGT, U/L | 10 - 60 | 78.90 (42.50,<br>179.75) | 46.70 (27.60,<br>70.00) | 26.30 (19.70,<br>43.50) | 27.10 (21.40,<br>41.70) | **0.002** | 0.058 | **<0.001** | **<0.001** | 0.486 |
| PLT, ×10<sup>9</sup>/L | 100 - 350 | 217.71 ± 75.23 | 256.65 ± 69.57 | 241.34 ± 58.84 | 250.96 ± 60.99 | 0.029 | 0.511 | 0.018 | 0.450 | 0.282 |
| TBiL, µmol/L | 1.7 - 17.1 | 9.74 ± 5.32 | 9.28 ± 4.35 | 9.32 ± 5.8 | 9.18 ± 6.03 | 0.769 | 0.680 | 0.326 | 0.869 | 0.374 |
| FPG, mmol/L | 3.9 - 6.1 | 7.06 ± 4.3 | 5.97 ± 2.99 | 5.37 ± 1.91 | 5.15 ± 1.43 | 0.404 | **0.004** | 0.016 | **0.004** | 0.519 |
| CRP, mg/L | < 8 | 3.51 (1.60, 5.73) | 2.35 (1.09,<br>5.25) | 1.81 (0.52, 3.71) | 1.37 (0.72,<br>2.93) | 0.276 | **0.004** | **<0.001** | 0.016 | **0.007** |
| TC, mmol/L | 2.84 - 5.68 | 4.81 ± 1.31 | 5.44 ± 1.26 | 5.31 ± 1.02 | 5.8 ± 1.24 | 0.910 | **<0.001** | **<0.001** | 0.012 | 0.108 |
| TG, mmol/L | 0.56 - 1.70 | 2.19 ± 1.13 | 2.32 ± 1.42 | 1.6 ± 0.85 | 1.83 ± 1.01 | 0.532 | 0.015 | **<0.001** | **<0.001** | 0.862 |
| HDL, mmol/L | 1.14 - 1.91 | 1.16 ± 0.33 | 1.29 ± 0.37 | 1.41 ± 0.35 | 1.43 ± 0.37 | 0.295 | 0.016 | **0.003** | **0.003** | 0.056 |
| LDL, mmol/L | 1 - 3 | 2.99 ± 1.00 | 3.37 ± 0.92 | 3.31 ± 0.78 | 3.66 ± 0.95 | 0.865 | 0.439 | 0.178 | **0.009** | 0.183 |
| UA, µmol/L | 90 - 360 | 351.37 ± 107.82 | 348.15 ± 91.19 | 330.4 ± 105.81 | 324.26 ± 85.09 | 0.272 | 0.981 | 0.430 | 0.021 | 0.543 |
| Urea, mmol/L | 1.8 – 7.1 | 6.16 ± 3.21 | 5.63 ± 1.54 | 6.05 ± 2.63 | 5.73 ± 1.44 | 0.211 | 0.127 | 0.014 | 0.581 | 0.903 |
| Comorbidities at baseline |  |  |  |  |  |  |  |  |  |  |
| Diabetes |  | 5 (6.41) | 4 (2.56) | 8 (10.26) | 20 (12.82) | 0.279 | 0.385 | 0.135 | **<0.001** | 0.569 |

|  |  |  |  |  |  |  |  |  |  |
| --- | --- | --- | --- | --- | --- | --- | --- | --- | --- |
| Hypertension | 9 (11.54) | 26 (16.67) | 19 (24.36) | 34 (21.79) | 0.400 | 0.060 | 0.084 | 0.315 | 0.783 |
| Digestive cancers | 3 (3.85) | 3 (1.92) | 2 (2.56) | 6 (3.85) | 0.661 | 1.000 | 1.000 | 1.000 | 0.899 |
| GIT/BT diseases | 10 (12.82) | 25 (16.03) | 19 (24.36) | 44 (28.21) | 0.650 | 0.100 | 0.014 | 0.014 | 0.639 |

Note: Data are presented as means ± SDs or frequencies (percentages). SD standard deviation, BMI body mass index; ALT, alanine transaminase; AST, aspartate transaminase; ALB, albumin; ALP, alkaline phosphatase; GGT, gamma-glutamyl transferase; PLT, platelet; TBI, total bilirubin; FPG, fasting plasma glucose; CRP, C-reactive protein; TC, total cholesterol; TG, triglycerides; HDL, high-density lipoprotein; LDL, low-density lipoprotein; UA, uric acid; MASH, metabolic dysfunction-associated steatohepatitis; MAS, metabolic dysfunction-associated steatosis, defined as MASLD without steatohepatitis; VH, viral hepatitis; NL, normal liver function control subjects. $P < 0.01$ (Bonferroni adjusted $\alpha$ ) is considered statistically significant. Significant $P$ -values are shown in bold italics.

Baseline liver function tests (ALT, AST, ALP and GGT) were significantly elevated in the MASH group compared to the normal liver control group at baseline levels, although ALP was still within the normal range for MASH cases (all *P* < α_adjust_, 0.01). No significant differences were observed between the two groups for ALB, PLT and TBiL levels (all *P* > 0.01). Significant differences in carbohydrate and lipid metabolism markers, including BMI, CRP, TC, TG, and HDL, were noted between the MASH and normal liver groups (all *P* < 0.01). Kidney function markers, such as urinary acid and urea, showed no significant differences between the MASH and normal liver groups (all *P* > 0.01) (**Table 1**).

Significant differences were identified in BMI, ALT, AST, FPG, TC, and TG, but not in other parameters in the MASH group when compared to the viral hepatitis group (**Table 1**). Taken together, these baseline characteristics suggest significantly elevated parameters in liver function in patients with MASH long before their corresponding disease diagnosis.

No significant differences were observed between the MASH and MAS groups in liver function and carbohydrate and lipid metabolism, except for the GGT baseline level. These baseline characteristics suggest that liver function and carbohydrate and lipid metabolism in MASH and MAS cases were similar at the baseline levels, indicating that these parameters are not sufficient to differentiate MASH cases from patients with MAS (**Table 1**).

### Expression levels of six proteomic biomarkers in MASH at baseline

The UK Biobank prospective cohort provided proteomic data to examine the baseline levels of six previously identified diagnostic biomarkers for MASH, MASLD, and liver cirrhosis. The mean time from the detection of biomarkers at baseline levels to the establishment of clinical diagnosis was over 10 years (**Fig. 1B**).

We compared the levels of six proteins among the four groups and found that the expression levels of circulating CDCP1, FABP4, FGF21, GDF15, IL-6, and THBS2 at baseline were highest in the MASH group (all *P* < 0.01), followed by a gradually decreasing trend in MAS, viral hepatitis, and normal liver control groups (**Table 2 and Fig. 2**).

**Table 2.** Levels of proteins in MASH and three control groups.

|  | Detected range in<br>blood by PEA | MASH<br>(n = 78) | MAS<br>(n = 156) | VH<br>(n = 78) | NL<br>(n = 156) |
| --- | --- | --- | --- | --- | --- |
| **CDCP1**, NPX | 1.0-4.0 | 0.91 ± 0.88 | 0.39 ± 0.74 | 0.36 ± 0.90 | 0.11 ± 0.62 |
| **FGF21**, NPX | 3.0-8.0 | 1.63 ± 1.65 | 1.18 ± 1.47 | -0.18 ± 1.42 | 0.02 ± 1.33 |
| **IL-6**, NPX | 2.0-6.0 | 1.19 ± 1.11 | 0.55 ± 0.79 | 0.30 ± 0.88 | 0.19 ± 0.97 |
| **FABP4**, NPX | 3.0-6.0 | 0.86 ± 0.92 | 0.54 ± 0.86 | 0.11 ± 0.82 | 0.03 ± 0.75 |
| **GDF15**, NPX | 3.0-5.0 | 1.15 ± 0.91 | 0.45 ± 0.71 | 0.37 ± 0.70 | 0.09 ± 0.51 |
| **THBS2**, NPX | 4.0-6.0 | 1.09 ± 1.06 | 0.37 ± 0.67 | 0.21 ± 0.61 | -0.01 ± 0.31 |

Note: NPX, normalized protein expression; PEA, proximity extension assay; MASH, metabolic dysfunction-associated steatohepatitis; MAS, metabolic dysfunction-associated steatosis, defined as MASLD without steatohepatitis; VH, viral hepatitis; NL, normal liver function control subjects; CDCP1, CUB domain-containing protein 1; FABP4, fatty acid binding protein 4; FGF21, fibroblast growth factor 21; GDF15, growth differentiation factor 15; IL-6, interleukin-6; THBS2, thrombospondin 2. The detected range of six circulating protein biomarkers in blood is obtained from The Human Protein Atlas (https://www.proteinatlas.org/).

**Fig. 2.**
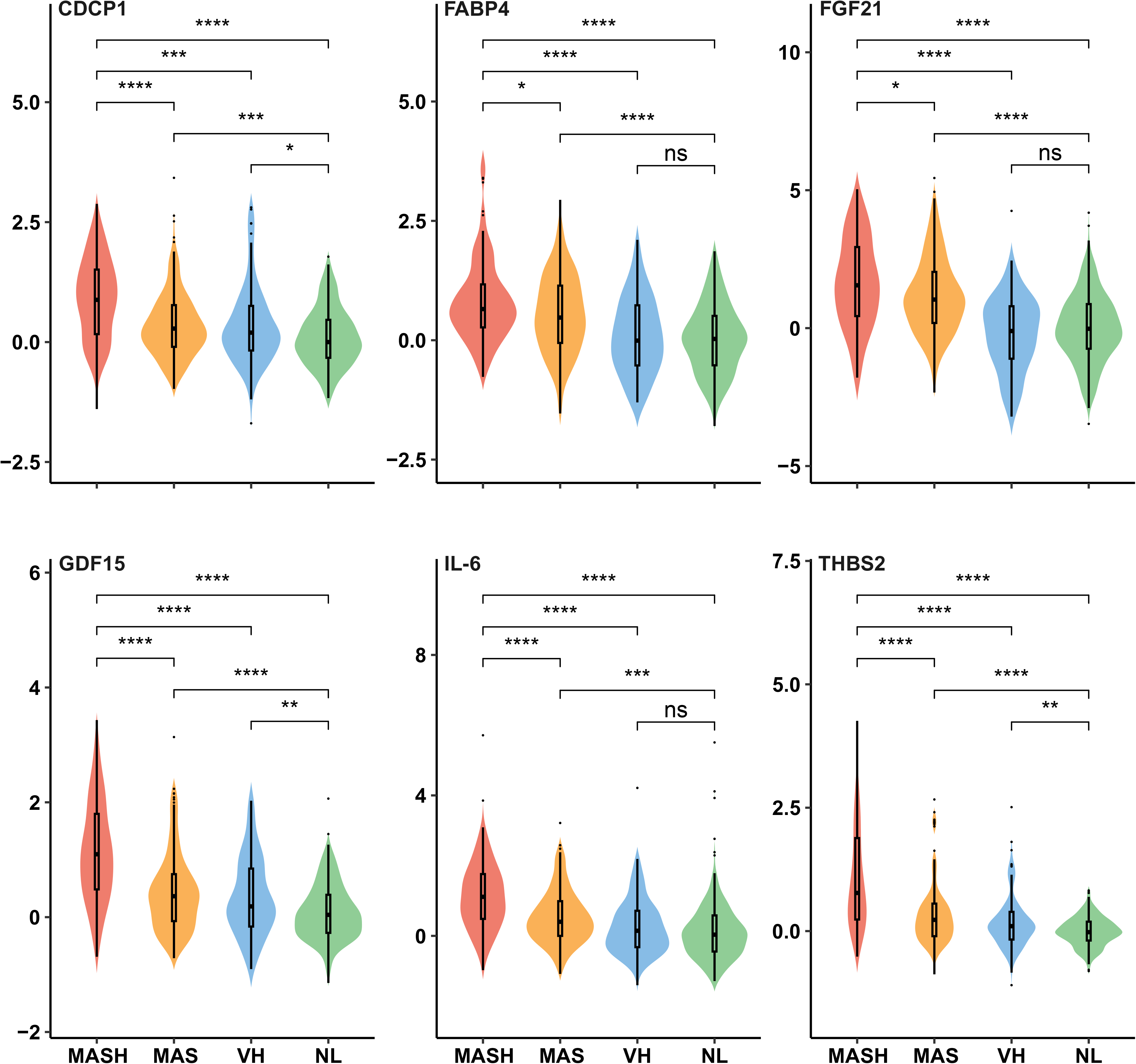
Violin plots comparing baseline levels of the six proteomic biomarkers among MASH, MAS, viral hepatitis and normal liver control groups. The expression levels of the six biomarkers were expressed as normalized protein expression (NPX) units. P value ranges were indicated with *s for comparisons among different groups. ns, non-significant; *; P < 0.05; **, P < 0.01; ***, P < 0.001; ****, P < 0.0001. *P* < 0.01 (Bonferroni adjusted α) is considered statistically significant. MASH, metabolic dysfunction-associated steatohepatitis. MAS, metabolic dysfunction-associated steatosis, defined as MASLD without steatohepatitis. VH: viral hepatitis. NL: normal liver function control subjects. CDCP1, CUB domain-containing protein 1; FABP4, fatty acid binding protein 4; FGF21, fibroblast growth factor 21; GDF15, growth differentiation factor 15; IL-6, interleukin-6; THBS2 thrombospondin 2.

Significantly higher levels of all six protein biomarkers were identified in the MASH group compared to the viral hepatitis and normal liver control groups (All *P* < 0.01). Except for FABP4 and FGF21, four biomarkers (CDCP1, GDF15, IL-6, and THBS2) were significantly higher in the MASH group compared to the MAS group at baseline levels (All *P* < 0.01). All six protein biomarkers were significantly higher in the MAS group compared to normal liver controls at baseline levels (All *P* < 0.01). There were no significant differences for CDCP1, FABP4, FGF21, and IL-6, between viral hepatitis group and normal liver controls (All *P* > 0.01, **Fig. 2**).

These findings highlight the potential of these six protein biomarkers, particularly GDF15, CDCP1, IL-6, and THBS2, to serve as early indicators of MASH. The elevated levels of these biomarkers in MASH patients long before the clinical diagnosis suggest their potential utility in the early detection and risk stratification of this disease.

### Association among baseline levels of the six circulating biomarkers and clinical characteristics

To determine the relationships among the six biomarkers, Spearman’s correlation coefficients were calculated and visualized in a correlation plot (**Fig. 3A**). The baseline levels of all biomarkers exhibited positive correlations with each other (*P* < 0.05). Notably, GDF15 and CDCP1 demonstrated the strongest positive correlation (*r* = 0.59, *P* < 0.05), suggesting a high degree of association between these two potential biomarkers. Conversely, FABP4 and CDCP1 showed the weakest positive correlation (*r* = 0.23, *P* < 0.05).

**Fig. 3.**
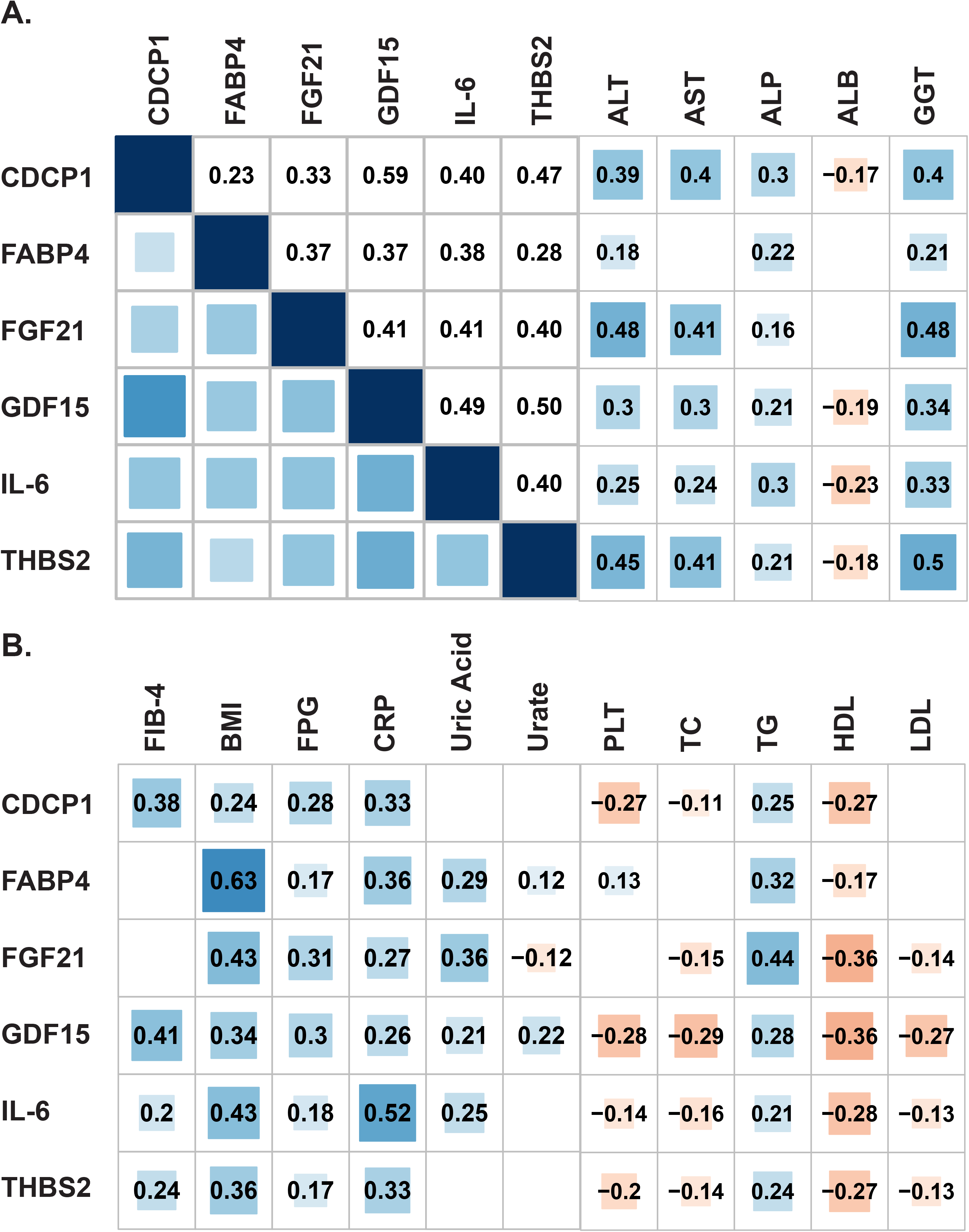
Spearman correlations among baseline levels of the six circulating biomarkers and clinical characteristics. Statistically significant correlations are shown with numbers, while the non-significant correlation coefficients are blank in the boxes. Positive correlations are represented by blue color and negative correlations are represented by red color. (A) Correlation plot of the six biomarkers and their relationship with liver function. (B) Correlation plot of the six biomarkers with other clinical factors. CDCP1, CUB domain-containing protein 1; FABP4, fatty acid binding protein 4; FGF21, fibroblast growth factor 21; GDF15, growth differentiation factor 15; IL-6, interleukin-6; THBS2, thrombospondin 2; ALT, alanine transaminase; AST, aspartate transaminase; ALB, albumin; ALP, alkaline phosphatase; GGT, gamma-glutamyl transferase; PLT, platelet; TBiL, total bilirubin; TC, total cholesterol; TG, triglycerides; HDL, high-density lipoprotein; LDL, low-density lipoprotein; UA, uric acid; CRP, C-reactive protein; and FPG, fasting plasma glucose.

We also calculated the correlation coefficients between protein biomarkers and liver function, kidney function, and metabolic tests. The six protein biomarkers displayed positive correlations with ALT, AST, ALP and GGT (**Fig. 3A**). CDCP1, FABP4, GDF15, IL-6, and THBS2 showed negative correlations with TBiL and/or ALB (**Fig. 3B**). Furthermore, the six biomarkers at baseline levels positively correlated with carbohydrate and lipid metabolism markers, including BMI, FPG, CRP, and TG, and negatively correlated with TC and HDL (**Fig. 3B**), highlighting their potential relevance to MASH pathology. Taken together, these correlation analyses underscore the interconnected nature of these biomarkers and their association with liver function and metabolic health.

### Risk effects of the six circulating protein biomarkers for MASH

Previous studies have identified the six biomarkers for diagnosis purpose based on blood samples obtained with biopsy-confirmed diagnoses. To determine the risk effects of these circulating biomarkers for MASH development prior to diagnosis, we calculated the HRs and 95% CIs using Cox regression. Three models were employed to adjust the clinical factors to remove potential confounding bias. Model 1 adjusted for one metabolic marker and two liver function markers, including BMI, ALT, and AST. Model 2 adjusted for one metabolic marker and all liver function markers, including BMI, ALT, AST, ALB, ALP, GGT, and PLT. Model 3 adjusted for all metabolic and liver function markers, including BMI, ALT, AST, ALB, ALP, GGT, PLT, TC, TG, HDL, and LDL.

Among the six biomarkers, GDF15 at baseline levels showed the highest HR for discriminating MASH from normal liver controls. After adjustment using model 1, high GDF15 baseline level was associated with a roughly 2.5-fold increased risk of MASH (HR = 2.49, 95% CI: 1.85-3.36, *P* < 0.001). Further adjustments using model 2 revealed that high GDF15 baseline level was associated with a 2.9-fold increased risk of MASH (HR = 2.89, 95% CI: 1.96-4.25, *P* < 0.001). After additional adjustments using model 3, the HR for GDF15 decreased slightly but remained highly significant (HR = 2.338, 95% CI: 1.53-3.54, *P* < 0.001, **Fig. 4A**). The elevated levels of the other five protein biomarkers were also associated with an increased the risk of MASH when compared to normal liver controls, though their risk effects were not as strong as GDF15 (**Fig. 4A**). Only two biomarkers, FGF21 and THBS2, were associated with increased risk of MAS development (**Fig. 4B**). None of the six circulating biomarkers at baseline levels were associated with increased risk of viral hepatitis (**Fig. 4C**). In conclusion, even after adjusting for clinical factors, GDF15 remained the most predictive biomarker for MASH, indicating its potential utility in early prediction and intervention strategies.

**Fig. 4.**
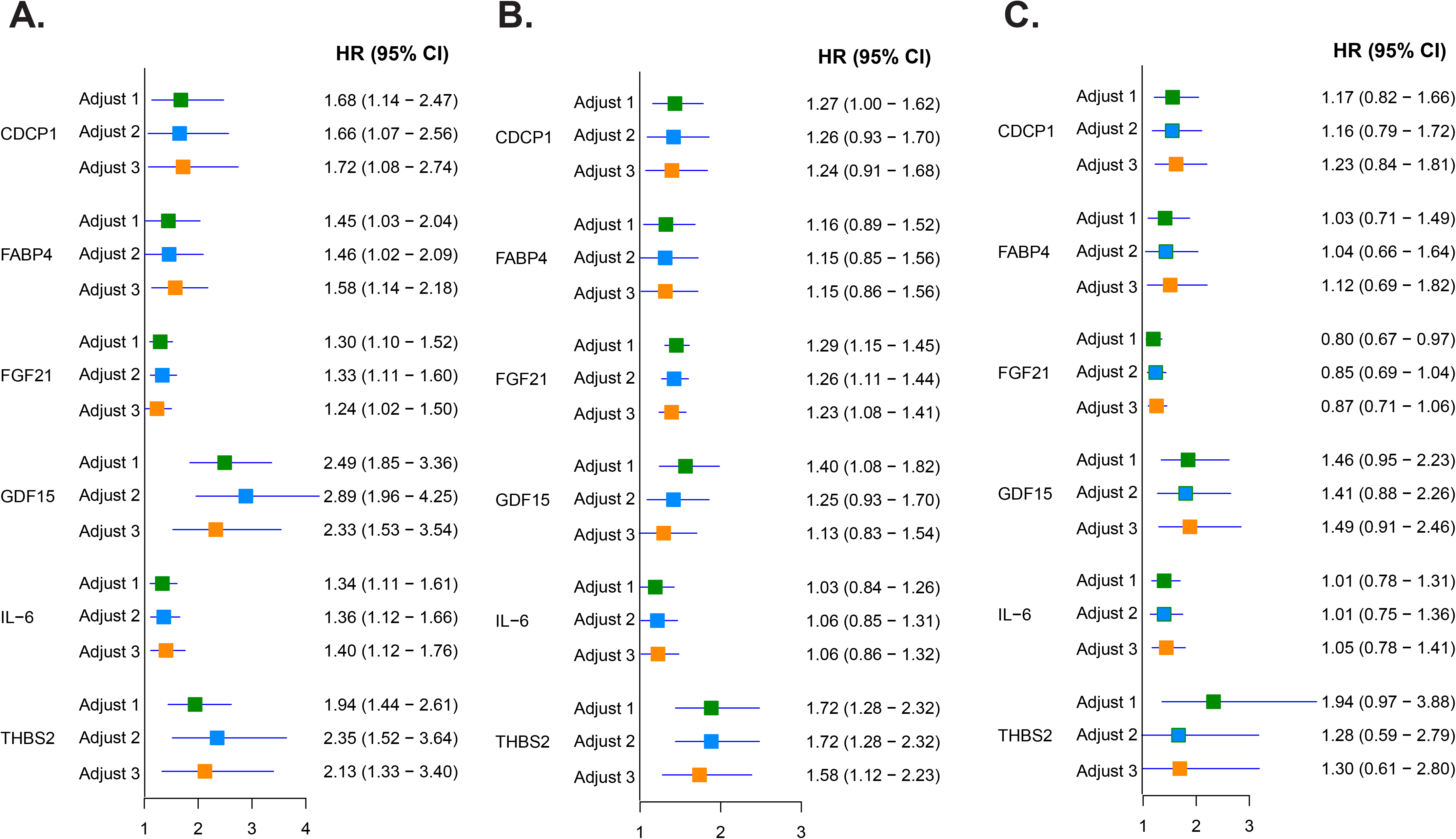
Forest plot of risk effects of the six circulating protein biomarkers for MASH, MAS and viral hepatitis development. HRs and 95% CIs were calculated using Cox regression. Three models were used to adjust the confounding factors with model 1 adjusted for BMI, ALT, and AST, model 2 adjusted for BMI, ALT, AST, ALB, ALP, GGT, and PLT, and model 3 adjusted for BMI, ALT, AST, ALB, ALP, GGT, PLT, TC, TG, HDL, and LDL. Shown are forest plots of HRs adjusted by three models for the six biomarkers in predicting MASH against NL (A), MAS against NL (B), and VH against NL (C). MASH, metabolic dysfunction-associated steatohepatitis; MAS, metabolic dysfunction-associated steatosis, defined as MASLD without steatohepatitis. VH: viral hepatitis. NL: normal liver function control subjects.

### Predictive performance of the six circulating biomarkers at baseline for subsequent MASH development

To evaluate the predictive performance of the six protein biomarkers for MASH development, time-dependent ROC curves were plotted individually. GDF15 at baseline level demonstrated the best performance for predicting MASH occurrence against normal liver controls. Specifically, the AUC for GDF15 at 5 years was 0.90 (95% CI: 0.82-0.97), and at 10 years, it was 0.86 (95% CI: 0.80-0.92) (**Fig. 5**).

**Fig. 5.**
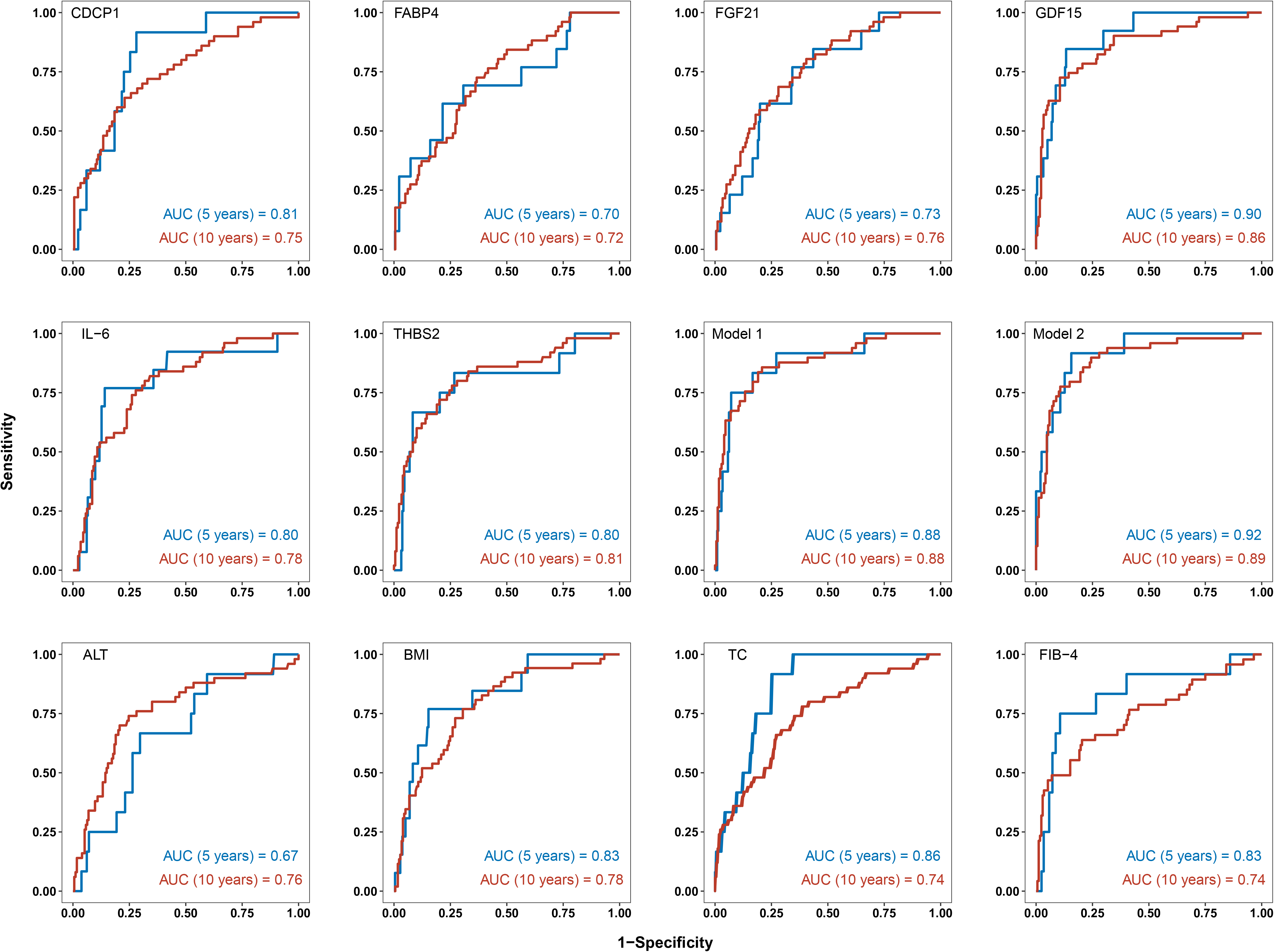
Time-dependent ROC curves of the six biomarkers, two combinatory models, and clinical factors for MASH prediction against normal liver controls. Shown are AUC values for MASH prediction against normal liver controls at 5 and 10 years of mean lag time from recruitment to diagnosis. Model 1 contains 4 biomarkers (GDF15, FGF21, IL-6 and THB2). Model 2 contains the 4 biomarkers from Model 1 and 3 clinical factors (BMI, ALT and TC). MASH, metabolic dysfunction-associated steatohepatitis.

Further analysis involved constructing predictive models based on the six protein biomarkers using Cox regression models. The best model was selected using the stepwise (both directions) Akaike’s Information Criterion (AIC) method. This process identified four key biomarkers (GDF15, FGF21, IL-6, and THBS2) for the final predictive model. The predictive performance of this model showed an AUC of 0.88 at both 5 years and 10 years (**Fig. 5**).

To enhance predictive accuracy further, a protein-clinical model was developed by incorporating clinical factors at baseline levels along with the protein biomarkers. This model included four key biomarkers (GDF15, FGF21, IL-6, and THBS2) and three clinical factors (BMI, ALT and TC), resulting in an AUC of 0.92 at 5 years and 0.89 at 10 years (**Fig. 5**). The predictive performances of the routinely clinical parameters, including ALT, BMI, TC and CRP, yielded AUCs lower than 0.9. (**Fig. 5**) Importantly, GDF15 also exhibited the best performance for predicting MASH occurrence against MAS at 5 and 10 years, with AUCs of 0.85 and 0.78, respectively. The protein model for MASH occurrence showed AUCs of 0.82 at 5 years and 0.77 at 10 years, while the protein-clinical model showed AUCs of 0.86 at 5 years and 0.80 at 10 years (**Fig. 6**). Furthermore, GDF15 at baseline level demonstrated strong predictive accuracy for distinguishing MASH from viral hepatitis, with AUCs of 0.84 at 5 and 0.77 at 10 years (**Fig. S1**).

**Fig. 6.**
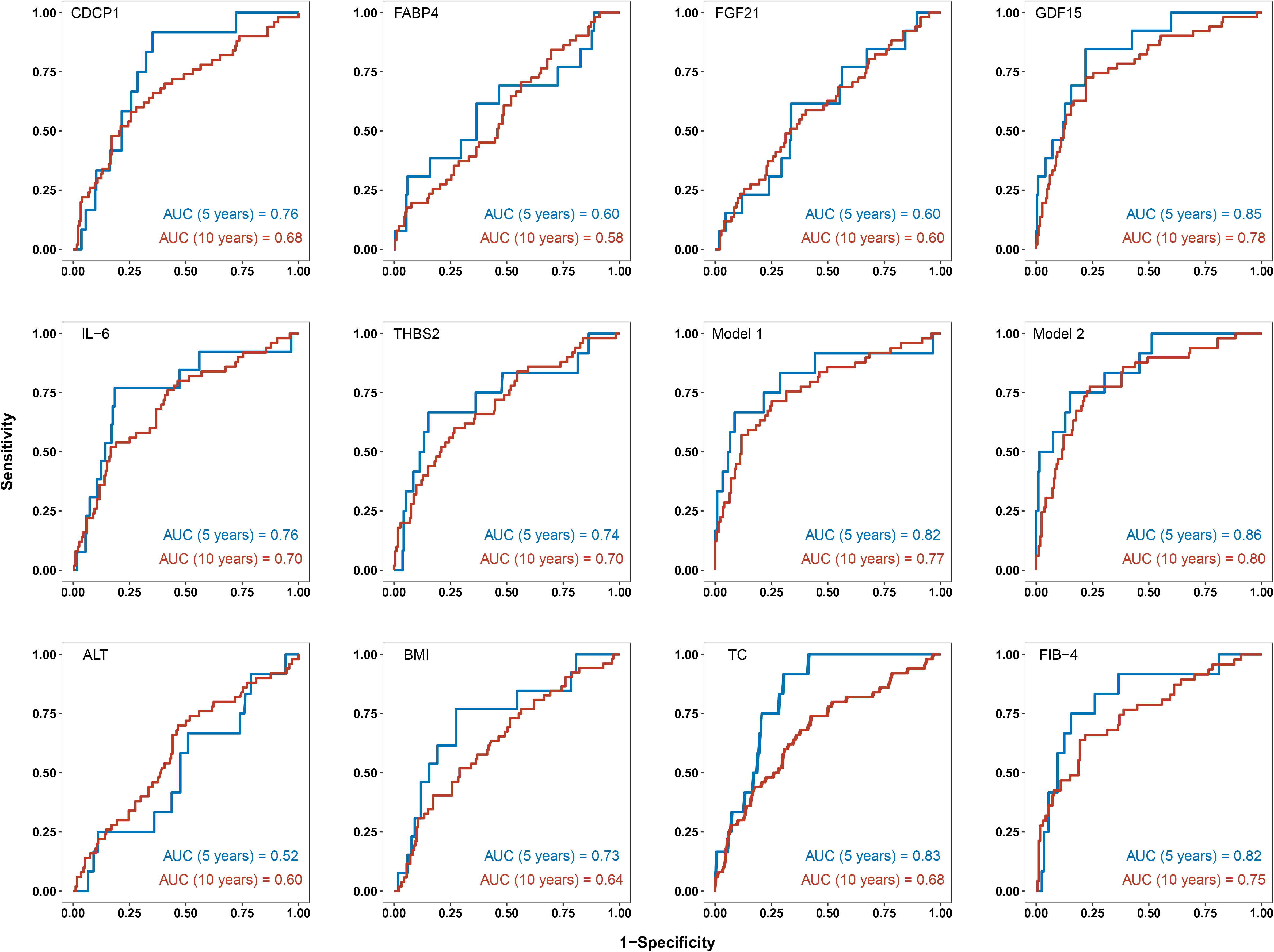
Time-dependent ROC curves of the six biomarkers, two combinatory models, and clinical factors for MASH prediction against MAS. Shown are AUC value for MASH prediction against MAS at 5 and 10 years of mean lag time from recruitment to diagnosis. Two combinatory biomarker panels were developed. Model 1 contains 4 biomarkers (GDF15, FGF21, IL-6 and THB2). Model 2 contains the 4 biomarkers from Model 1 and 3 clinical factors (BMI, ALT and TC). MASH, metabolic dysfunction-associated steatohepatitis; MAS, metabolic dysfunction-associated steatosis, defined as MASLD without steatohepatitis.

The predictive performance of these protein biomarkers for MAS were also evaluated individually, but none yielded AUCs higher than 0.7 for predicting MAS occurrence against normal liver controls at 5 or 10 years (**Fig. S2**). Predictive models for MAS, based on the protein biomarkers and clinical factors, also showed relatively poor performance. The protein model for MAS had AUCs of 0.619 at 5 years and 0.719 at 10 years, while the protein-clinical model had AUCs of 0.672 at 5 years and 0.734 at 10 years (**Fig. S2**). Additionally, the performance of the six individual biomarkers and developed predictive models for viral hepatitis was inferior to their performance for predicting MASH and MAS, highlighting the excellent specificity of these biomarkers for MASH prediction (**Fig. S3**).

In conclusion, GDF15 at baseline demonstrated superior predictive accuracy for MASH development, outperforming the other five biomarkers. The combination of protein biomarkers and clinical factors further enhanced predictive performance, with GDF15 playing a pivotal role. These findings underscore the specificity and utility of these biomarkers, particularly GDF15, in early MASH prediction. However, their effectiveness in predicting MAS and viral hepatitis was notably lower, emphasizing their specificity for MASH.

## Discussion

The lack of reliable, non-invasive biomarkers for early prediction and risk stratification of patients with MASH poses a significant challenge for timely intervention. In this study, we investigated the predictive accuracy of six previously identified diagnostic biomarkers including CDCP1, FABP4, FGF21, GDF15, IL-6, and THBS2 for predicting MASH development using proteomic data from a prospective cohort in the UK Biobank. Our investigation identified GDF15 at baseline levels as the most effective predictive biomarker for stratifying patients at high risk of developing MASH compared to those who later developed MAS, virial hepatitis, or individuals with normal liver function. Furthermore, we developed two models with high sensitivities and specificities for predicting MASH based on these circulating protein biomarkers. Our findings suggest that proteomics biomarkers hold promising clinical applications in the prediction and prevention of MASH.

GDF15, also known as macrophage inhibitory cytokine-1, belongs to the transforming growth factor β (TGF-β) superfamily and plays a crucial role in regulating TGF-β signalling.(23) It is closely involved in infection, fibrosis, and apoptosis pathways in response to tissue damage or disease progression.(16) In the context of MASH, liver fibrosis is a significant hallmark, where hepatic stellate cells (HSCs) play a pivotal role. TGF-β and related signals can accelerate HSC activation, promoting their proliferation, migration, contraction, and secretion of extracellular matrix proteins, thereby contributing to liver fibrogenesis.(24) Like other TGF-β superfamily cytokines, GDF15 plays an important role in pro-fibrotic effect in liver fibrogenesis.(25) A multicentre transcriptomic study demonstrated that hepatic GDF15 expression was positively associated with MASLD severity, and GDF15 expression was significantly higher in patients with advanced liver fibrosis.(26) Taken together, GDF15 may promote liver fibrogenesis in MASH progression, and the circulating level of GDF15 could be elevated long before disease diagnosis of MASH.

Clinically, the detected range of GDF15 in blood of large cohorts of old adults (>70 year old) and patients with chronic obstructive pulmonary disease by immunoassay method is about 1.1-1.4 µg/L,(27–29) and by proximity extension assay method is about 3.0-5.0 normalized protein expression (NPX, **Table 2**).(30) Our study found that the mean levels of GDF15 in MASH, MAS, and normal liver control groups is 1.63, 1.18 and 0.02 NPX, respectively, well below the above reported 3.0-5.0 NPX value. Furthermore, a previous study found that as diagnostic biomarkers, the mean levels of GDF15 by immunoassay in MASH patients, MAS patients and non-MASLD controls is 1.15 µg/L 0.97 µg/L, and 0.62 µg/L, respectively.(17) The mean levels of GDF15 in none/mild fibrosis (F0-1), moderate (F2), and severe fibrosis (F3-4) groups is 0.78 µg/L, 0.93 µg/L, and 1.50 µg/L, respectively.(17) A previous case-control study detected serum GDF15 using immunoassay and found that GDF15 is the most important single biomarker for the diagnosis of MASLD patients with moderate-to-severe fibrosis (F2-4 fibrosis).(31) The serum level of GDF15 is significantly higher in MASLD patients with advanced liver fibrosis, and it can distinguish F2-4 MASLD patients with a sensitivity of 0.56, a specificity of 0.87, and an AUC of 0.75.(31) The diagnostic cut-off value based on Youden index is 1.19 µg/L.(31) Our study found that GDF15 at baseline levels at 1.63 NPX is significantly increased in MASH patients, and it can predict MASH occurrence from different control groups with high sensitivity and specificity. In contrast, none of the elevated clinical factors at baseline levels can predict MASH development with high specificity. Comparing to serving as a diagnostic biomarker for MASH and MASLD, our findings suggest that GDF15 at baseline level could be used as a predictive biomarker for MASH long before its diagnosis. In future study, it will be important to measure the GDF15 at baseline using immunoassay to quantitate the concentration of GDF15.

Currently, many clinical diagnostic models and blood-based biomarkers have been developed as alternatives for the non-invasive diagnosis of patients with MASH.(9) Several these non-invasive diagnostic biomarkers may also predict adverse clinical outcomes, such as liver-related mortality, cardiovascular events, HCC, or liver transplantation.(32, 33) However, no highly sensitive and specific tests are available to predict MASH long before the disease diagnosis. The lack of MASH predictive biomarkers may be due to the slow progression from simple hepatic steatosis to MASH, and only a part of patients with simple hepatic steatosis can progress to MASH. Therefore, developing predictive biomarkers for MASH needs a long-term follow-up cohort with large sample size. The UK Biobank, with multi-omics data and long-term follow-up of large size participants, presents an invaluable resource for identifying accurate predictive biomarkers for MASH.

Our study, utilizing Olink-based proteome profiling, uniquely identifies GDF15 as a robust circulating protein biomarker for early MASH prediction, demonstrating its predictive efficacy with a mean lag time exceeding 10 years. While acknowledging limitations such as sample size and the need for further multi-centered validation, our findings suggest GDF15 and GDF15-based models as promising tools for predictive diagnosis and population risk stratification in MASH.

In conclusion, GDF15 emerges as a promising biomarker for MASH prediction, surpassing other studied protein biomarkers. Its incorporation into easy-to-implement predictive models could potentially reduce the need for invasive procedures like liver biopsies, benefiting both patients and healthcare systems alike. Continued research and validation efforts are essential to fully harness the predictive potential of GDF15 in clinical practice.

## Abbreviations

AIC: Akaike’s information criterion
ALB: albumin
ALP: alkaline phosphatase
ALT: alanine transaminase
AST: aspartate transaminase
AUC: area under the curve
CDCP1: CUB domain-containing protein 1
CI: confidence interval
CRP: C-reactive protein
FABP4: fatty acid binding protein 4
FGF21: fibroblast growth factor 21
FPG: fasting plasma glucose
GDF15: growth differentiation factor 15
GGT: gamma-glutamyl transferase
HCC: hepatocellular carcinoma
HDL: high-density lipoprotein
HR: hazard ratio
HSCs: hepatic stellate cells
LDL: low-density lipoprotein
ICD-10: International Classification of Diseases 10th revisionIL-6, interleukin-6
MAS: metabolic dysfunction-associated steatosis
MASH: metabolic dysfunction-associated steatohepatitis
MASLD: metabolic dysfunction-associated steatosis liver disease
NAFLD: non-alcoholic fatty liver disease
NL: normal liver function control subjects
NPX: normalized protein expression
PLT: platelet
ROC: receiver operating characteristic
SD: standard deviations
TBiL: total bilirubin
TGF-β: transforming growth factor β
TC: total cholesterol
TG: triglycerides
THBS2: thrombospondin 2
UA: uric acid
VH: viral hepatitis.

## Author contributions

HY, JJ, YWH, and YK: conceptualization; HW and YK: methodology; HW and YS: software; XX: validation; HW: formal analysis; XX and YS: investigation; HW: data curation; HW: writing-original draft preparation; YWH and YK: writing-review and editing; HW: visualization; JJ, and YK: supervision; HY and YK: funding acquisition. All authors have read and agreed to the published version of the manuscript.

## Acknowledgments

The authors acknowledge the participants and their families who contributed to make this study possible.

## Data availability statement

The UK Biobank data can be accessed through the UKBB portal (http://www.ukbiobank.ac.uk).

## Supplementary data

The supplementary methods and data to this article can be found online at

