## Supplementary Methods and Figures for "Identification of Growth Differentiation Factor-15 as An Early Predictive Biomarker for Metabolic Dysfunction-Associated Steatohepatitis: A Nested Case-control Study of UK Biobank Proteomic Data"

**Figure S2.** Time-dependent ROC curves of the six biomarkers, two combinatory models, and clinical factors for MASH prediction against VH…… ........................................................................................................................ 7

**Figure S3.** Time-dependent ROC curves of the six biomarkers, two combinatory models, and clinical factors for MAS prediction against normal liver controls... .......................................................................................................... 8

**Figure S4.** Time-dependent ROC curves of the six biomarkers, two combinatory models, and clinical factors for VH prediction against normal liver controls…… .................................................................................................... 9

### Supplementary Methods

#### *Study design*

To validate proteomic biomarkers within the UK Biobank, a nested case-control study was conducted. This approach involves selecting cases and controls from the population of an existing cohort, offering several advantages over traditional case-control studies. Firstly, this method allows for the selection of cases and controls from the same population, which enhances internal validity by ensuring that both groups are drawn from the same underlying cohort and demographic characteristics. This is crucial in minimizing biases that can arise from differences in population characteristics. Secondly, nested case-control studies often employ matching techniques to pair cases with controls based on factors such as age, sex, and other relevant variables. This matching helps to minimize potential confounding variables, thereby making the analysis of biomarkers more robust and reliable.

#### *UK Biobank study population*

The UK Biobank is a comprehensive prospective cohort study aimed at improving the prevention, diagnosis, and treatment of various serious illnesses. It enrolled over 500,000 participants aged 37-73 years from the general population, achieving a response rate of 5.5%.[1] Between 2006 and 2010, participants attended one of 22 assessment centres across Scotland, England, and Wales.[2] During these visits, participants completed touch-screen questionnaires, underwent physical measurements, and provided blood, urine, and saliva samples at baseline.

#### *Biochemical Measurements*

Routine biochemical tests were conducted to measure of the levels of several biomarkers, including alanine transaminase (ALT), aspartate transaminase (AST), albumin (ALB), alkaline phosphatase (ALP), gamma-glutamyl transferase (GGT), platelet (PLT), total bilirubin (TBiL), total cholesterol (TC), triglycerides (TG), high-density lipoprotein (HDL), low-density lipoprotein (LDL), uric acid (UA), urea, C-reactive protein (CRP), and fasting plasma glucose (FPG). These measurements were obtained from the UK Biobank and provide a comprehensive biochemical profile for each participant.

#### *Olink proteomics and candidate proteomic biomarkers data*

The UK Biobank Pharma Proteomics Project (UKB-PPP) conducted a large-scale plasma proteome study using the Olink Explore 1536 Proteomics platform. This study analyzed 53,014 plasma samples from individual UK Biobank participant-visits. The Olink platform utilizes Proximity Extension Assay (PEA) technology, which allows for the quantification of 2,939 protein analytes. These analytes span various panels including inflammation, cancer, cardiometabolic, and neurological markers.

In this project, PEA assays were implemented to measure protein expression levels, and extensive data pre-processing and normalization procedures were employed, as detailed in previous publications.[3] The resulting proteomic data is presented in log2 normalized protein expression (NPX) units, which standardizes the data for analysis and comparison across samples.

This study specifically focuses on utilizing baseline proteomic data from the UK Biobank cohort. Among the wealth of protein analytes measured, the NPX data of six candidate protein biomarkers were extracted for analysis: CDCP1, FABP4, FGF21, GDF15, IL-6, and THBS2. These biomarkers were selected based on their diagnostic relevance in distinguishing between various liver disease states, including MASH and related conditions. The use of the Olink Explore 1536 platform in conjunction with the UK Biobank provides a robust framework for investigating proteomic signatures associated with different health conditions.

***References***

[1] Collins R. What makes UK Biobank special? Lancet. 2012;379(9822): 1173-1174.

[2] Palmer LJ. UK Biobank: bank on it. Lancet. 2007;369(9578): 1980-1982.

[3] Sun BB, Chiou J, Traylor M, et al. Plasma proteomic associations with genetics and health in the UK Biobank. Nature. 2023;622(7982): 329-338.

Table S1. Exclusion criteria for diagnosis of other liver diseases

| **Disease** | **ICD-10** |
|---|---|
| Alcoholic liver disease | K70 |
| Toxic liver disease | K71 |
| Hepatic failure, not elsewhere classified | K72 |
| Chronic hepatitis, not elsewhere classified | K73 |
| Fibrosis and cirrhosis of liver | K74 |
| Other inflammatory liver diseases | K75 |
| Other diseases of liver | K76 |
| Liver disorders classified elsewhere | K77 |
| Malignant neoplasm of liver and intrahepatic bile ducts | C22 |
| Unspecified viral hepatitis | B19 |

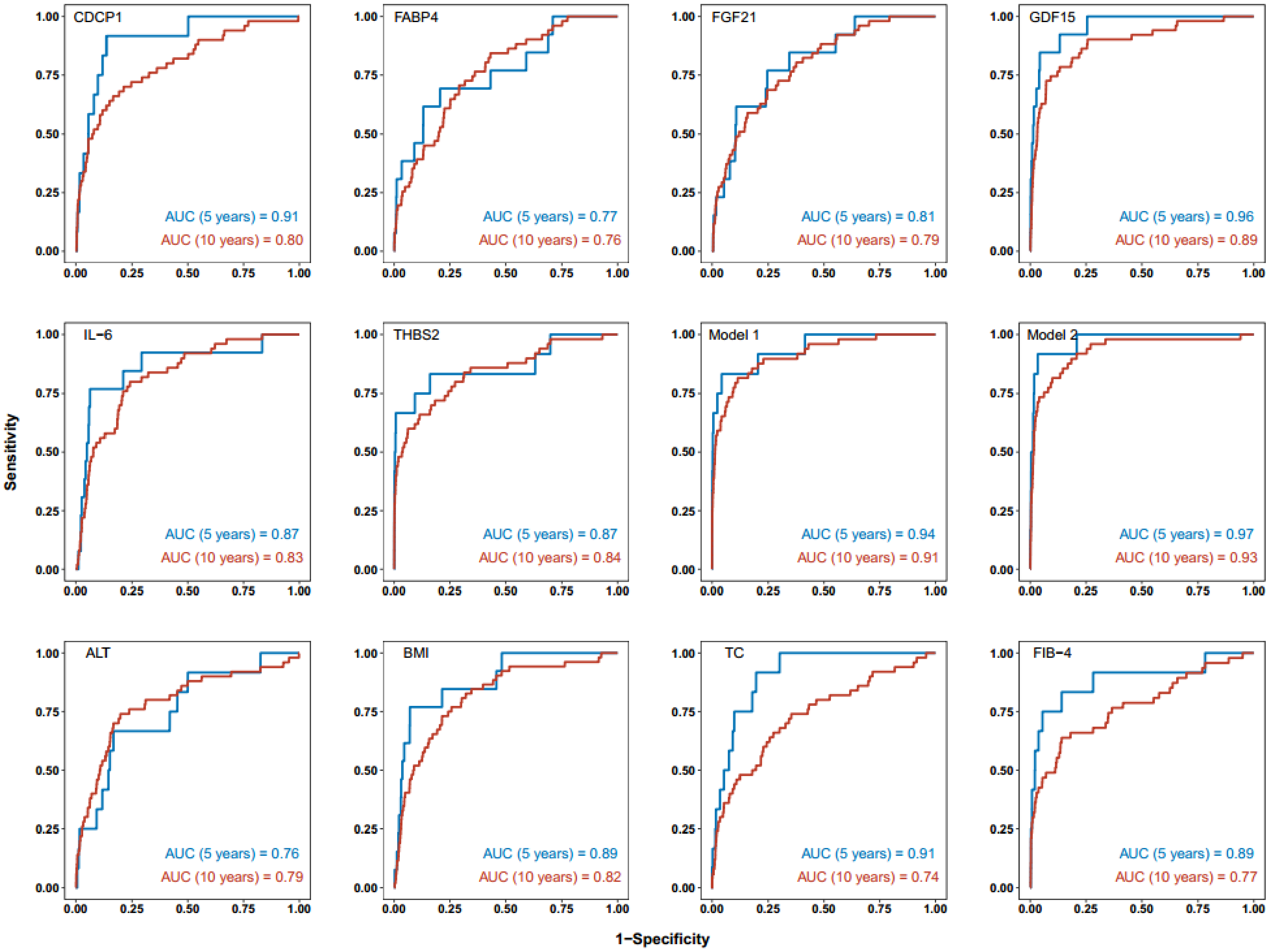

**Figure S1. Time-dependent ROC curves of the six biomarkers, two combinatory models, and clinical factors for MASH prediction against normal liver controls in general population.** Shown are AUC values for MASH prediction against normal liver controls in general population at 5 and 10 years of mean lag time from recruitment to diagnosis. Model 1 contains 4 biomarkers (GDF15, FGF21, IL-6 and THB2). Model 2 contains the 4 biomarkers from Model 1 and 3 clinical factors (BMI, ALT and TC). MASH, metabolic dysfunction-associated steatohepatitis.

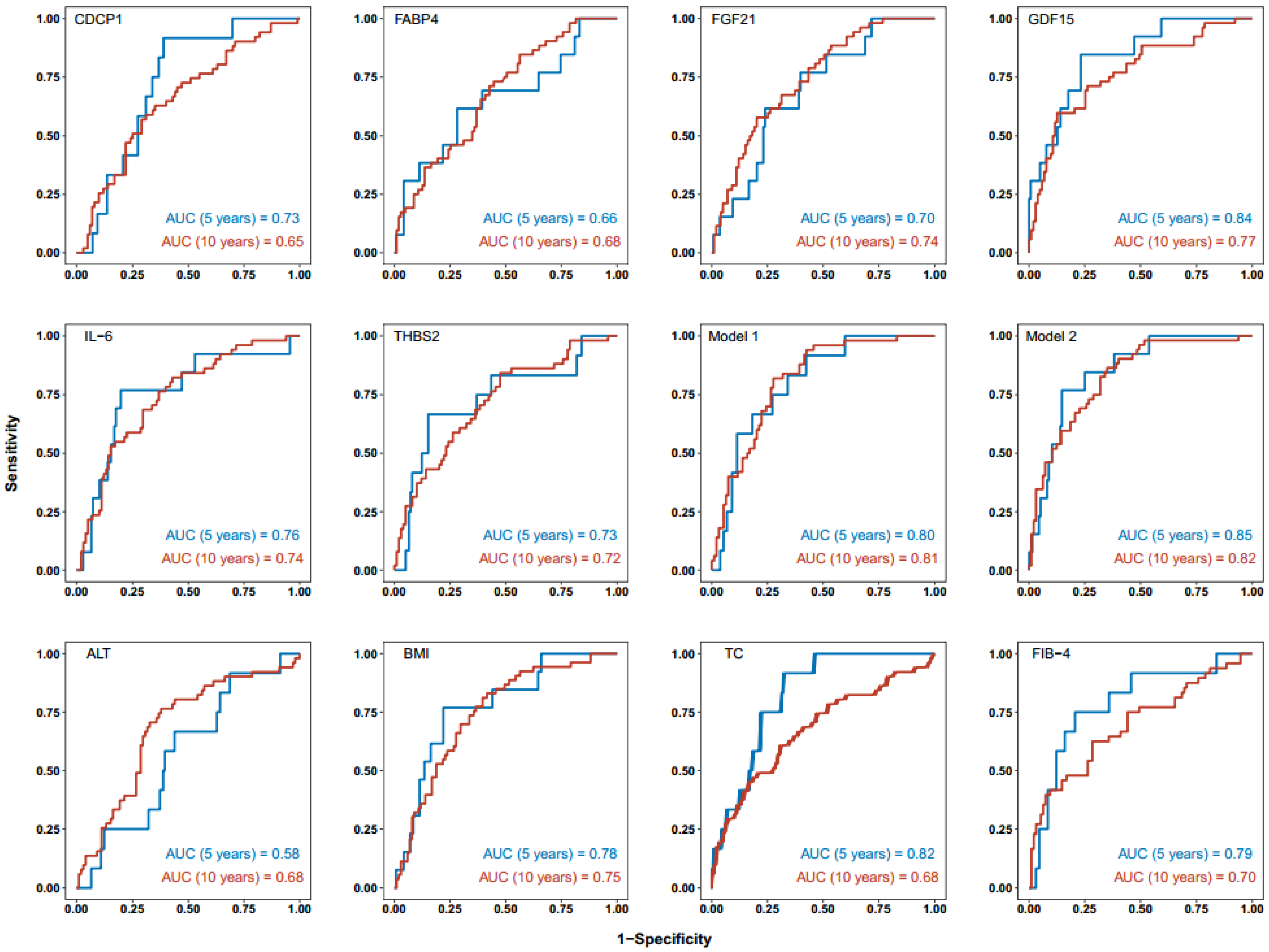

**Figure S2. Time-dependent ROC curves of the six biomarkers, two combinatory models, and clinical factors for MASH prediction against VH.** Shown are AUC value for MASH prediction against viral hepatitis at 5 and 10 years of mean lag time from recruitment to diagnosis. Two combinatory biomarker panels were developed. Model 1 contains 4 biomarkers (GDF15, FGF21, IL-6 and THB2). Model 2 contains the 4 biomarkers from Model 1 and 3 clinical factors (BMI, ALT and TC). MASH, metabolic dysfunction-associated steatohepatitis; VH, viral hepatitis.

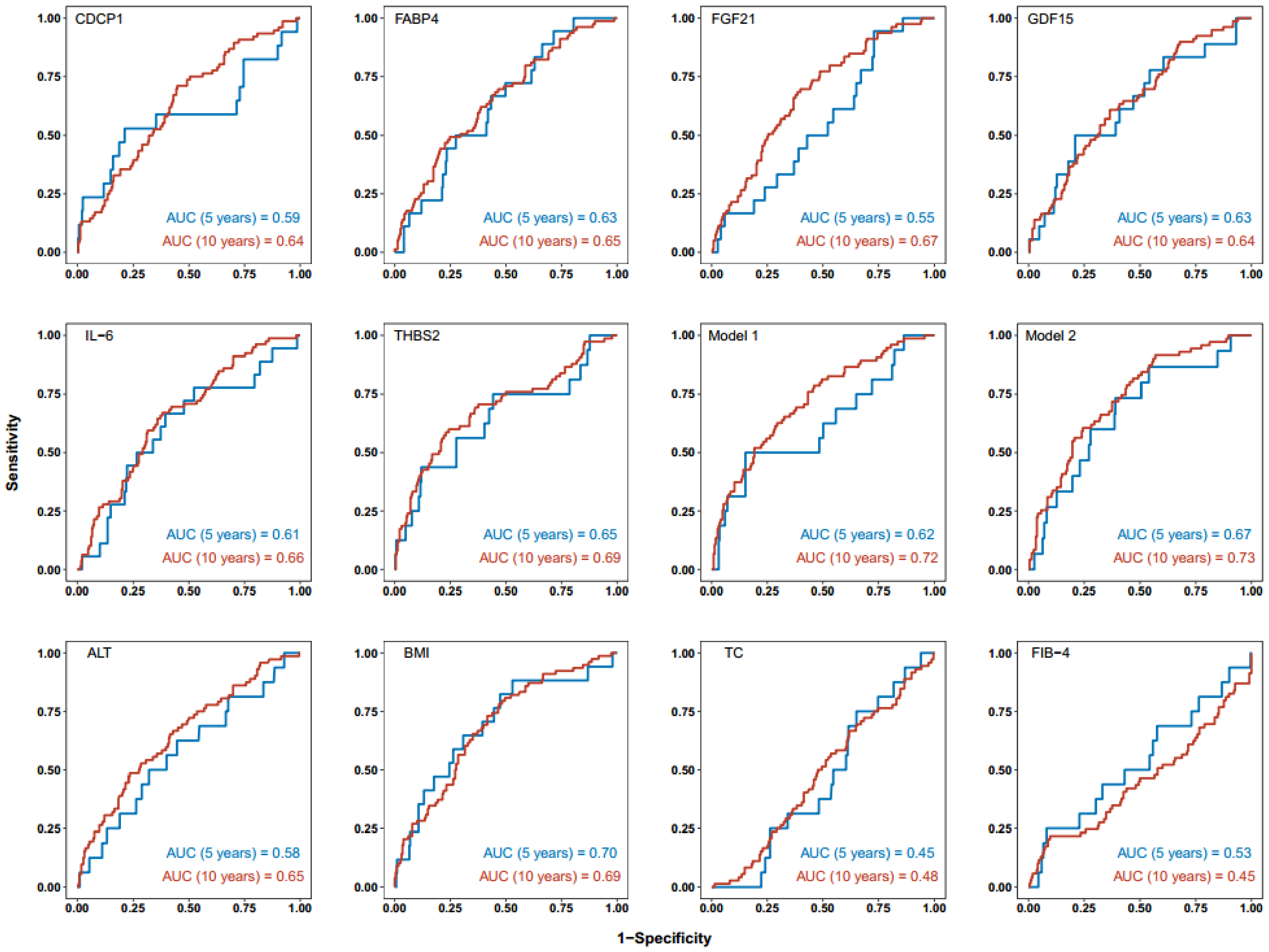

**Figure S3. Time-dependent ROC curves of the six biomarkers, two combinatory models, and clinical factors for MAS prediction against normal liver controls.** Shown are AUC values for MAS prediction against normal liver controls at 5 and 10 years of mean lag time from recruitment to diagnosis. Two combinatory biomarker panels were developed. Model 1 contains 4 biomarkers (GDF15, FGF21, IL-6 and THB2). Model 2 contains the 4 biomarkers from Model 1 and 3 clinical factors (BMI, ALT and TC). MAS, metabolic dysfunction-associated steatosis, defined as MASLD without steatohepatitis.

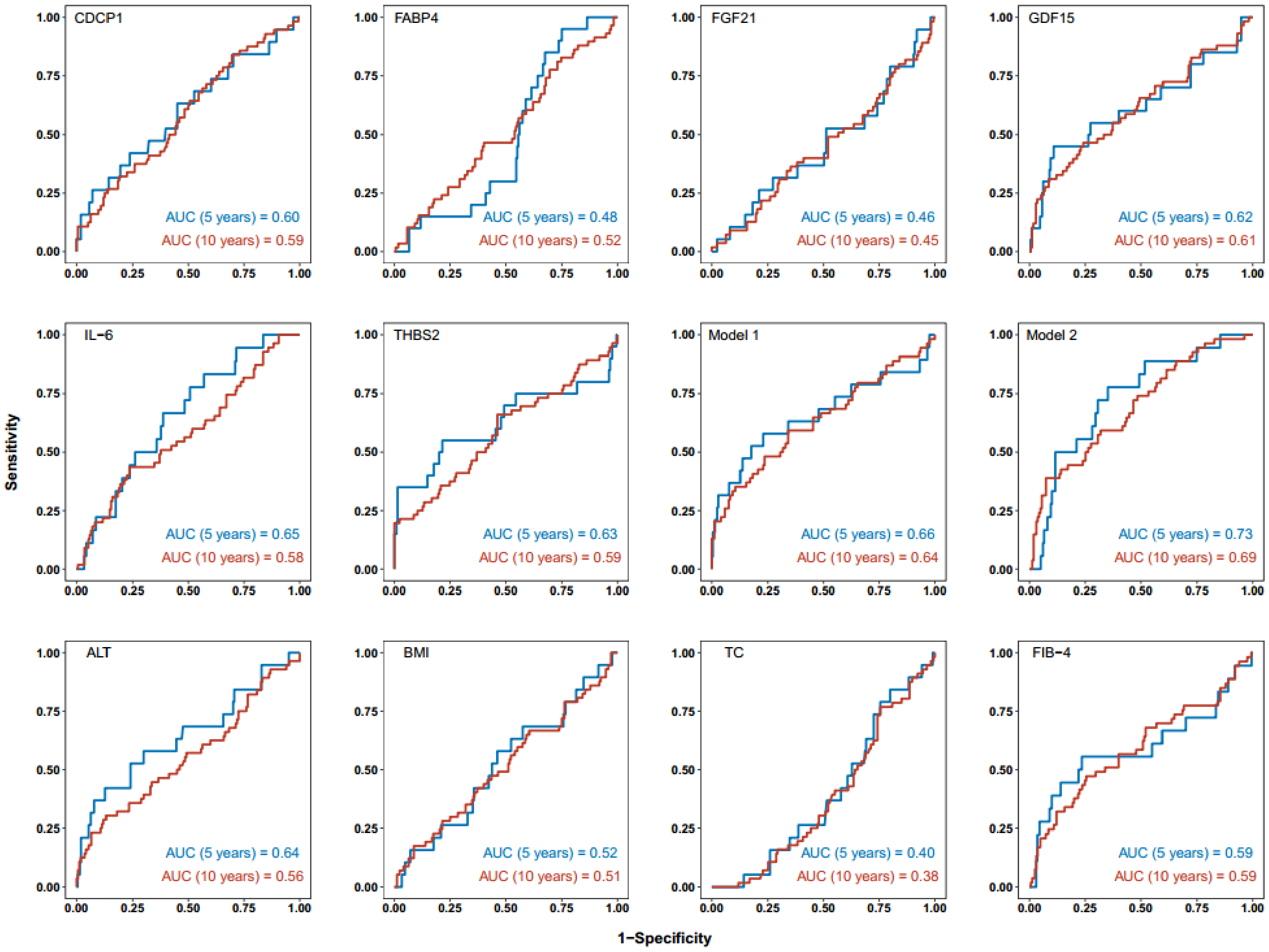

**Figure S4. Time-dependent ROC curves of the six biomarkers, two combinatory models, and clinical factors for VH prediction against normal liver controls.** Shown are AUC values for VH prediction against normal liver controls at 5 and 10 years of mean lag time from recruitment to diagnosis. Two combinatory biomarker panels were developed. Model 1 contains 4 biomarkers (GDF15, FGF21, IL-6 and THB2). Model 2 contains the 4 biomarkers from Model 1 and 3 clinical factors (BMI, ALT and TC). VH, viral hepatitis.
